# The MoT3 assay does not distinguish between *Magnaporthe oryzae* wheat and rice blast isolates from Bangladesh

**DOI:** 10.1101/345215

**Authors:** Dipali Rani Gupta, Claudia Sarai Reyes Avila, Joe Win, Darren M. Soares, Lauren S. Ryder, Daniel Croll, Pallab Bhattacharjee, Md. Shaid Hossain, Nur Uddin Mahmud, Md. Shabab Mehbub, Musrat Zahan Surovy, Nicholas J. Talbot, Sophien Kamoun, M. Tofazzal Islam

## Abstract

The blast fungus *Magnaporthe oryzae* is comprised of lineages that exhibit varying degrees of specificity on about 50 grass hosts, including rice, wheat and barley. Reliable diagnostic tools are essential given that the pathogen has a propensity to jump to new hosts and spread to new geographic regions. Of particular concern is wheat blast, which has suddenly appeared in Bangladesh in 2016 before spreading to neighboring India. In these Asian countries, wheat blast strains are now co-occurring with the destructive rice blast pathogen raising the possibility of genetic exchange between these destructive pathogens. We assessed the recently described MoT3 diagnostic assay and found that it did not distinguish between wheat and rice blast isolates from Bangladesh. The assay is based on primers matching the WB12 sequence corresponding to a fragment of the *M. oryzae* MGG_02337 gene annotated as a short chain dehydrogenase. These primers could not reliably distinguish between wheat and rice blast isolates from Bangladesh based on DNA amplification experiments performed in separate laboratories in Bangladesh and in the UK. In addition, comparative genomics of the WB12 sequence revealed a complex underlying genetic structure with related sequences across *M. oryzae* strains and in both rice and wheat blast isolates. We, therefore, caution against the indiscriminate use of this assay to identify wheat blast.

## INTRODUCTION

Outbreaks caused by fungal diseases have increased in frequency and are a chronic threat to global food security (Fisher et al. 2012). A prime case is blast, a disease caused by the ascomycete fungus *Magnaporthe oryzae* (syn. *Pyricularia oryzae*), which is best known as the most destructive disease of rice. However, in addition to rice, *M. oryzae* can also infect other cereal crops such as wheat, barley, oat and millet destroying food supply that could feed hundreds of millions of people (Pennisi 2010; Fisher et al. 2012; Liu et al. 2014). Increased global trade, climate change, and the propensity of this pathogen to occasionally jump from one grass host to another, have resulted in increased incidence of blast diseases. For example, only a few decades ago blast was not known to affect wheat, a main staple crop critical to ensuring global food security. But in 1985, blast disease on wheat was first reported in Paraná State, Brazil (Igarashi et al. 1986). It has since spread throughout Brazil and subsequently moved into neighboring South American countries where it is now a major threat to wheat production (Goulart et al. 1992; Goulart et al. 2007; Kohli et al. 2011). Currently, wheat blast affects as many as 3 million hectares of cultivated wheat, seriously limiting the potential for wheat production in the vast grasslands region of South America.

More recently, in February 2016, wheat blast was detected for the first time in Asia, following reports of a severe outbreak in Bangladesh (Islam et al. 2016).The outbreak was particularly destructive, affecting ~16% of the cultivated wheat area in Bangladesh and with yield losses reaching up to 100% (Islam et al. 2016). In 2017, the pathogen spread to neighboring West Bengal region of India and hundreds of hectares were cleared by burning to limit the accumulation and spread of the pathogen according to local press reports. An open source population genomics project led by Islam *et al*. (2016) revealed that the Bangladeshi wheat blast outbreak was caused by a South American genotype of *M. oryzae*, and therefore most likely introduced from South America (OpenWheatBlast http://wheatblast.net). The 2016 Bangladeshi epidemic vividly illustrates the clear and present danger caused by blast diseases in an era of global trade. It is especially worrisome because blast could spread further to other wheat producing areas in South Asia, such as India and Pakistan, thus threatening food security across South Asia.

Until recently, the genetic structure of *M. oryzae* remained somewhat unclear. Whole genome sequence analyses of 76 isolates from 12 different grass hosts confirmed the status of *M. oryzae* as a single species and revealed genetic exchanges between the different host-specific lineages (Gladieux et al. 2018). Overall, *M. oryzae* lineages exhibit low levels of nucleotide polymorphisms within single copy genes, with π ranging from 7.75e^-04^ to 1.24e^-03^ in the various host-specific lineages of this species. This is one order of magnitude lower than divergence between *M. oryzae* and its closest relatives *Magnaporthe grisea* and *Magnaporthe pennisetigena* (Gladieux et al. 2018). Nonetheless, the different lineages of *M. oryzae* exhibit significant genome plasticity probably due to the activity of transposable elements and selection for optimal disease effector repertoires (Yoshida et al. 2016). Notably, effector genes often exhibit presence/absence polymorphisms, and particular effector genes can be associated with a given lineage probably contributing to the specialization of individual genotypes to certain hosts (Inoue et al. 2017).

Given the propensity of *M. oryzae* to jump hosts and spread to new geographic regions, reliable and cost-effective diagnostic tools are needed to monitor this pandemic pathogen at the genotype level. This is not necessarily a straightforward problem considering the relatively low genetic diversity and potential for gene flow between lineages of *M. oryzae*. Pieck *et al*. (2017) recently reported a polymerase chain reaction (PCR) assay diagnostic for wheat blast based on a single molecular marker named MoT3. The assay is based on primers matching the WB12 sequence corresponding to a fragment of the *M. oryzae* MGG_02337 gene annotated as a short chain dehydrogenase. Here, we report that these primers could not reliably distinguish between wheat and rice blast isolates from Bangladesh based on DNA amplification experiments performed in separate laboratories in Bangladesh and in the UK. In addition, comparative genomics of the WB12 sequence revealed a complex underlying genetic structure with WB12 related sequences occurring across *M. oryzae* strains and in both rice and wheat blast isolates. We, therefore, caution against the indiscriminate use of this assay to identify wheat blast.

## RESULTS AND DISCUSSION

### MoT3 PCR assays yield similar sized bands with *Magnaporthe oryzae* wheat and rice blast isolates

To evaluate the MoT3 assay, we first performed PCR using MoT3 primers in the laboratory of MTI in Bangladesh. We used as template genomic DNA of *M. oryzae* isolates collected from wheat (BTMP 13-1, BTMP 13-2, BTJP 4-5, and BTJP4-6) and rice plants (RB-3b, RB-9d, RB-11b, and RB-11d) that showed blast disease symptoms during the 2016 and 2017 epidemics in Bangladesh. We used PoT2 transposon primers as positive control for presence of *M. oryzae* genomic DNA. We found that both primer pairs amplified expected size amplicons (361 and 389 bp for MoT3 and PoT2, respectively) from all genomic DNA from *M. oryzae* isolates (Fig. 1).

**Figure 1.**
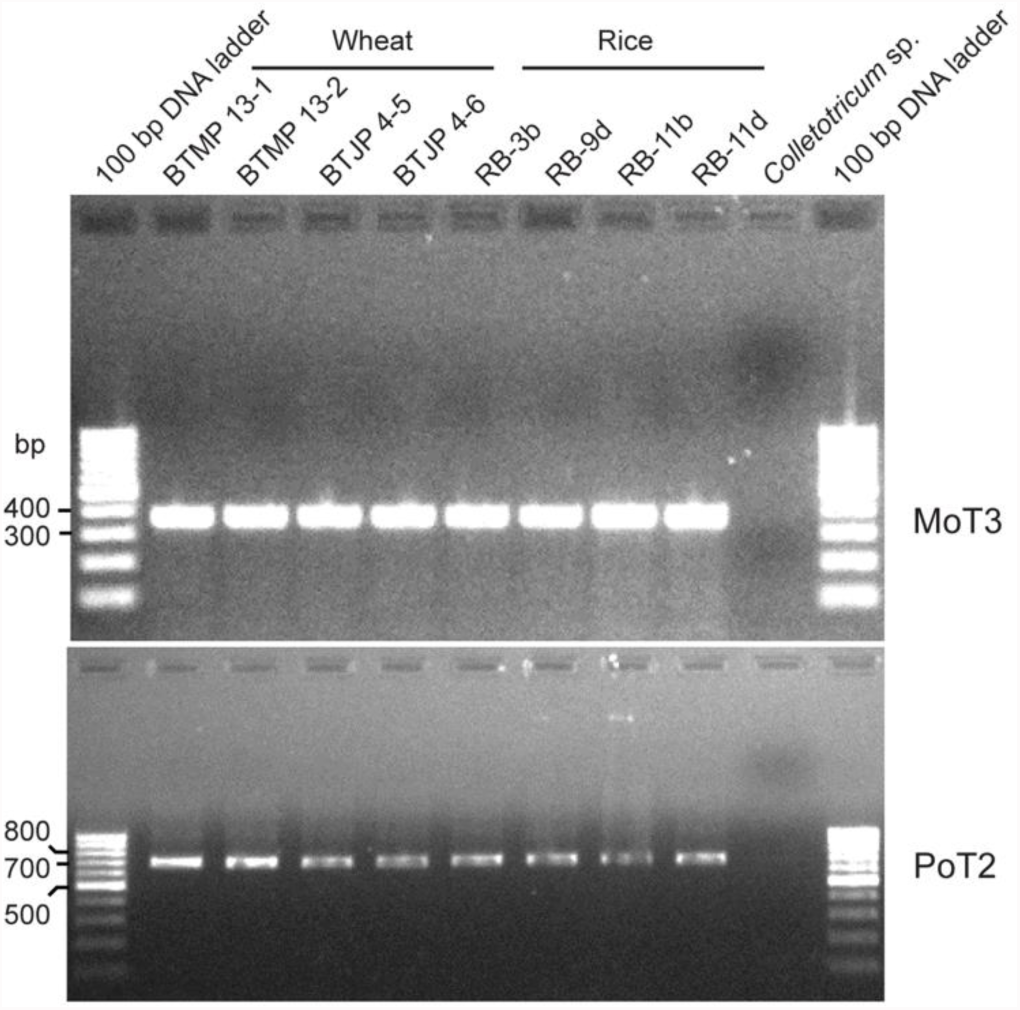
Amplification of wheat and rice blast isolates by MoT-3 primers and PoT2 primers.

We independently repeated the MoT3 assays in The Sainsbury Laboratory, UK, with wheat blast isolate BTJP 4-1 and three rice blast isolates (RB9c, RB11a, and RB13-1c) from Bangladesh, along with the reference wheat isolate BR32 from Brazil and rice isolate INA168 from Japan (Fig. 2).

**Fig. 2.**
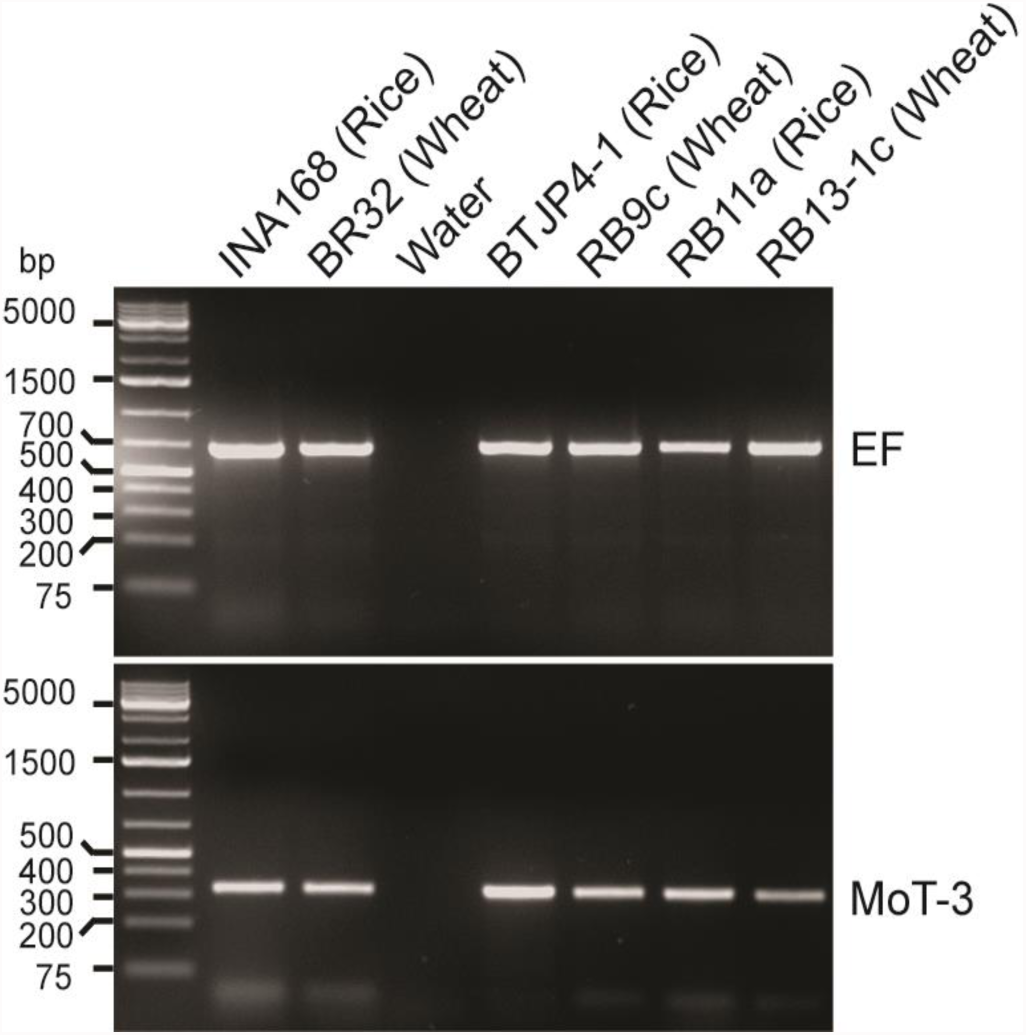
Amplification of WB12 region from rice and wheat-infecting *M. oryzae* using MoT-3 primers and elongation factor primers. Amplicons from MoT3 primers and elongation factor primers are labelled with MoT-3 and EF, respectively. GeneRuler Plus (Thermo-Fisher) DNA ladder was used as standard markers.

We used elongation factor (EF) primers as positive control. Both primer pairs amplified amplicons of the expected size (361 and 722 bp for Mot3 and EF, respectively) with all tested isolates (Fig. 2)

### MoT3 PCR amplifications yield diverse sequences

To further explore the PCR assays shown in Fig. 2, we sequenced the amplicons. We then constructed a multiple sequence alignment using the amplicon sequences (excluding the primer sequences) along with genomic sequences from WB12-like regions of several *M. oryzae* isolates identified by BLASTN search of genome sequences available from The *Magnaporthe* GEMO Database website (http://genome.jouy.inra.fr/gemo/) (Chiapello et al. 2015) (Fig. 3A). Amplicon sequences from *M. oryzae* from Bangladesh rice blast isolates (RB9c, RB11a, and RB13-1c) showed that they bear different WB12 sequences despite the fact that MoT3 primers were able to produce amplicons with expected size from these isolates (Fig. 3). The Bangladesh wheat blast isolate shows high sequence identity to WB12 sequence itself, whereas the Bangladeshi rice isolates has high degree of similarity with genome sequence of reference rice isolates (FR13, GY11, PH14-rn, TH16,) and the one isolate from *Eleusine* (CD156). The WB12-like sequence of *M. oryzae* wheat isolate BR32 from Brazil was more similar to those from rice isolates than to the wheat isolates indicating that the distribution of the different WB12 sequences is not strictly correlated with host or origin (Fig. 3B).

**Fig. 3.**
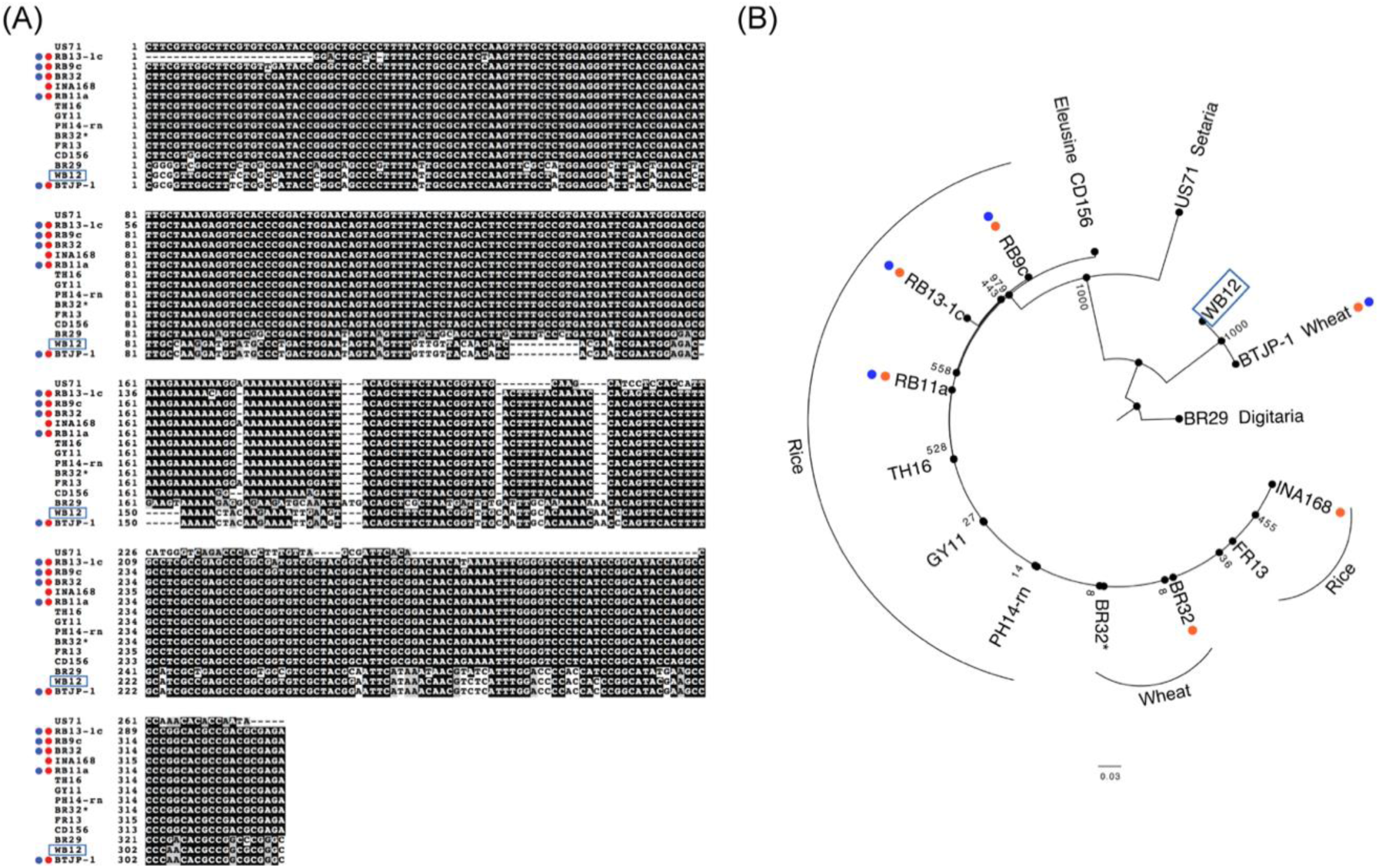
MoT3 amplicon sequences. (A) Multiple sequence alignment of WB12 sequence (boxed in blue), MoT3 amplicon sequences (indicated by red dots), and WB12-like genomic regions of several *M. oryzae* isolates. Sequence conservation for each site is indicated by box-shading the nucleotide with either black (>50%) or grey (“conservative mutation”). (B) A gene tree based on sequences of WB12 region in a selection of *M. oryzae* isolates. Red dots = Amplicon sequences; Blue dots = Bangladeshi isolates; the rest of the sequences were extracted from the genomic sequences.

**Fig. 4.**
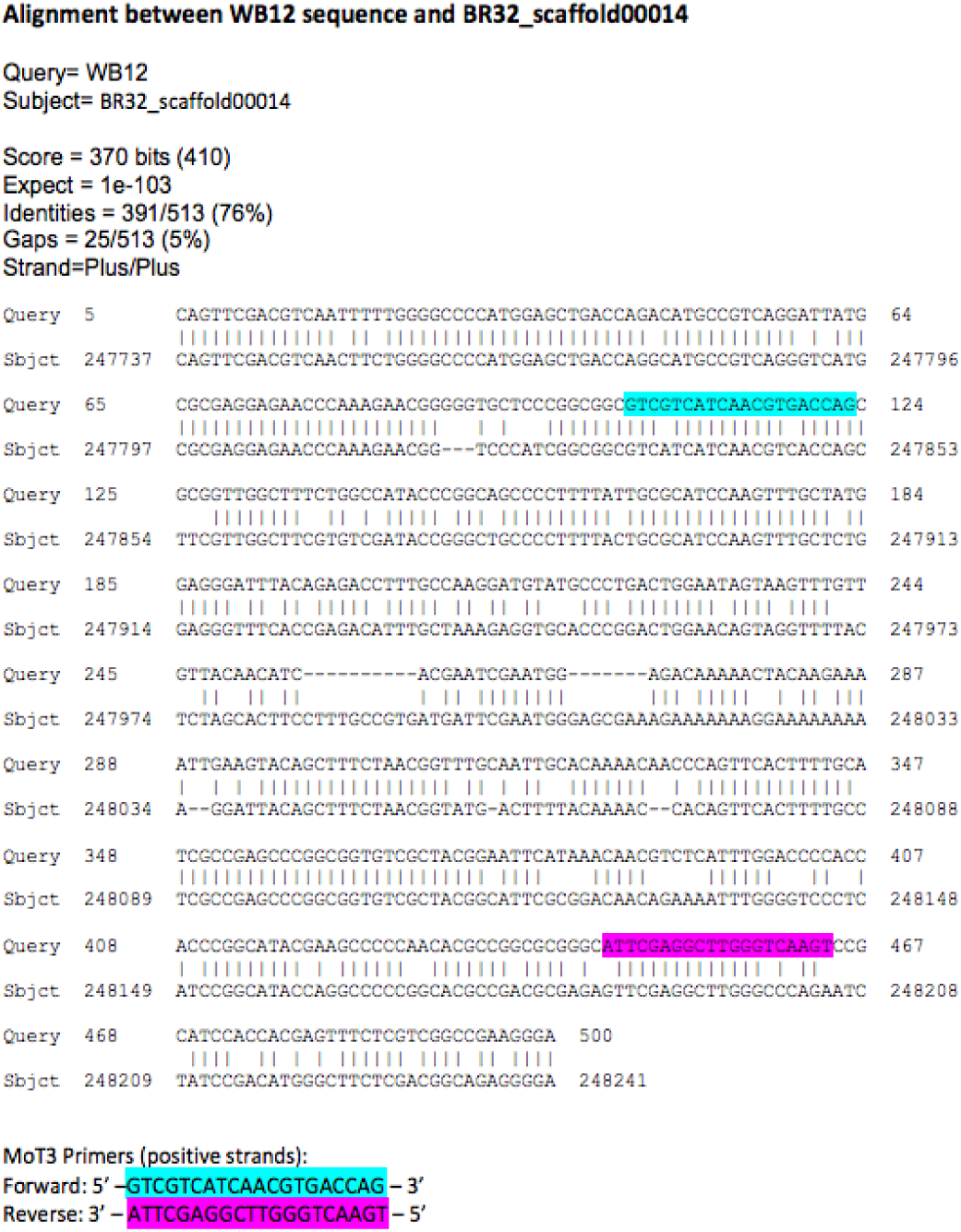
Alignment of second WB12 region in BR32 against WB12 with MoT3 primers highlighted: Forward primer in cyan; Reverse primer in magenta.

### Comparative genomics analyse s reveal that WB12-paralogous sequences can serve as templates for MoT3 primers

To assess the diversity and distribution of the WB12 sequence within populations of *M. oryzae*, we searched the genome sequences of 67 *M. oryzae* isolates from various host plants available in Genbank using BLAST v.2.2.26. We found significant hits with >98% identity across at least 400 bp in the majority of isolates from the wheat-infecting lineage, including *Bromus* isolate P29, which was previously grouped as a member of a wheat-infecting lineage (Suppl. Table 1). However, three wheat isolates, notably BR32, did not yield a BLAST match to WB12 (Suppl. Table 1). We conclude that the WB12 sequence is not present in all wheat-infecting isolates.

BLASTN searches also produced sequences related to WB12 (we refer to as WB12-like) in several contigs of the assembled genomes of the 67 *M. oryzae* isolates. The first set of WB12-like sequence had 70-98% identity to the WB12 sequence over a 400 bp matched region, and was present in 65 isolates except in US71 and GrF52 from *Setaria* spp. The second set of WB12-like had 60-98% identity over 100-400 bp of the matching region, and was only found in US71 and GrF52. We conclude that WB12 region is specific to a subset of wheat-infecting isolates of *M. oryzae* and missing in other lineages (Suppl. Table 1). In contrast, the WB12-like sequences are present in almost all wheat and non-wheat isolates (Suppl. Table 2).

We checked the degree to which the MoT3 primers could match the WB12-like sequences. MoT3 primer sequences can be aligned with high identity to the WB12-like sequence. For example, MoT3 primer sequences can be aligned to WB12-like sequence of *M. oryzae* wheat isolate BR32 (Suppl. Fig 1). Although there were a few mismatches between primers and BR32 genome sequence, these sequence differences apparently do not interfere with PCR amplification under the conditions we described in the methods.

## CONCLUSIONS

Our findings lead us to caution against the indiscriminate use of the MoT3 assay to identify wheat blast. The MoT3 primers can under certain amplification conditions yield positive amplicons with the great majority of *M. oryzae* isolates independently of their host of origin.

## METHODS

### MoT3 assay

To confirm that MoT3 primers amplify genomic DNA of wheat blast fungi, we used genomic DNA extracted from one *M. oryzae* isolate from Bangladeshi wheat blast (BTJP4-1) and one from Brazilian wheat blast (BR32) as templates. To test the specificity of MoT3 primers, we also included genomic DNA from three *M. oryzae* isolates from Bangladeshi rice blast (RB9c, RB11a, and RB13-1c) and one from Japan (INA168).

PCR assays with MoT3 primers (MoT3F 5’-GTCGTCATCAACGTGACCAG-3’ and MoT3R 5’-ACTTGACCCAAGCCTCGAAT-3’) (Pieck et al. 2017), and elongation factor-1α primers (forward 5’-CTYGGTGTTAGGCAGCTCA-3’ and reverse 5’-GAAMTTGCAGGCRATGTGGG-3’) (Castroagudin et al. 2016) were performed following the methods described in Pieck *et al.* (2017). Briefly, 50 µl PCR reactions containing 1x DreamTaq Green buffer (Thermo-Fisher), 0.2 mM dNTPs, 200 nM each primers, 100 ng template genomic DNA, and one unit of DreamTaq polymerase (Thermo-Fisher) were set up and amplification was performed using C1000 Thermal Cycler (Bio-Rad) with the following program: initial denaturation at 94 °C for 60 s; 30 cycles of 94 °C for 30 s, 62 °C for 30 s, 72 °C for 90 s; and extension at 72 °C for 120 s.

PCR products (10 µl each) were run on 1.5% agarose gel and stained with ethidium bromide. The rest of the PCR products were purified with PCR purification Kit (Qiagen) and sent for sequencing from both strands at GATC Biotech. DNA sequence contigs were assembled from sequencing chromatograms using Sequencher (Genecodes) for each amplicon and exported in fasta format for further analysis.

Sequences from each amplicon contig were aligned together with genomic regions from several *M. oryzae* isolates corresponding to the target sequence WB12 described in Pieck *et al.* (2017). These regions were identified by BLASTN in the genomic sequences of *M. oryzae* isolates from rice (FR13, GY11, PH14-rn, TH16), *Eleusine indica* (CD156), *Setaria italica* (US71) and wheat (BR32), as well as one *M. grisea* isolate (BR29) from *Digitalis sanguinalis* (downloaded from http://genome.jouy.inra.fr/gemo/, (Chiapello et al. 2015)). Primer sequences were removed from the alignment and a gene tree was constructed using clustalw2 neighbour joining algorithm and bootstrapped 1000 times.

### WB12 sequence similarity analyses

We used BLASTN program from BLAST suite version 2.2.26 (Altschul et al. 1997) to search WB12 sequence (Pieck et al. 2017) in the genome assemblies of 67 *M. oryzae* isolates, originating from various hosts and described in Genbank.

## ACKNOWLEDGEMENTS

This work was funded by the World Bank, the Biotechnology Biological Sciences Research Council (BBSRC), the European Research Council (ERC) and the Gatsby Charitable Foundation.

## SUPLEMENTARY MATERIAL

Table S1. BLASTN searches of WB12 against *M. oryzae* genomes.

Table S2. Highlights from BLASTN (less stringent) searches of WB12-like regions against 67 genomic sequences.

